# Mapping cell migrations and fates from a gastruloid model to the human primitive streak

**DOI:** 10.1101/616227

**Authors:** I. Martyn, E.D. Siggia, A.H. Brivanlou

## Abstract

Although fate maps of early gastrula embryos exist for nearly all model organisms, a fate map of the gastrulating human embryo remains elusive. Here we use human gastruloids to piece together part of a rudimentary fate map of the human primitive streak (PS). This is possible because stimulation with differing levels of BMP, WNT, and NODAL leads to self-organization of gastruloids into large and homogenous different subpopulations of endoderm and mesoderm, and comparative parallel analysis of these gastruloids, together with the fate map of the mouse embryo, allows the organization of these subpopulations along an anterior-posterior axis. We also developed a novel cell tracking technique that allowed the detection of robust fate-dependent cell migrations in our gastruloids comparable to those found in the mouse embryo. Taken together, our gastruloid derived fate map and recording of cell migrations provides a first coarse view of the embryonic human PS.

---

During amniote gastrulation a symmetric sheet of identical cells rapidly transforms itself into a multi-layered structure with distinct cell fates and an anterior-posterior axis^1^. This process commences with the initiation of the primitive streak, a transient structure which begins on the posterior edge and grows towards the center. As the streak grows, cells migrate through it and give rise to the different endodermal and mesodermal lineages of the future body plan. Specification of these different cell types depends on the position and time at which a cell transits through the streak.

A first step toward understanding early gastrulation in any organism is to track the developmental path of each progenitor cell and learn what structures and lineages they contribute to at later times. This information can be represented graphically in a so-called “fate map” where cell location and fate can be labelled and marked. When combined with perturbation studies that disrupt normal development, these maps can be useful in pinpointing the nature of the disruption and inferring cause and effect.

Although such fate maps have been completed for most model organisms^2–6^, no such fate map exists for human. This is because of ethical reasons prohibiting culturing human embryos beyond 14-days *ex vivo*, the time when the primitive streak first appears^7–9^. A map of the gastrulating human embryo, however, could be incredibly useful not just for comparison to model organisms and for understanding development in general, but for the practical use to efficiently guiding directed differentiation strategies of human embryonic stem cells (hESCs) into endoderm or mesoderm cell subtypes or even into a whole organ.

Given the restrictions alternative strategies have been pursued. Recently, we have proposed an alternative “gastruloid” approach to study early human gastrulation that is robust and amenable to single-cell quantification^10,11^. We have shown that when grown in epiblast-like geometrically confined disks, hESCs respond to BMP4 by differentiating and self-organizing into concentric rings of embryonic germ layers: with ectoderm in the center, extra-embryonic tissue at the edge, and mesoderm and endoderm in between. We have also shown an evolutionary conserved BMP→WNT→NODAL signalling hierarchy is responsible for this patterning, with WNT being necessary and sufficient to induce the primitive streak and NODAL biasing the ratio of mesodermal versus endodermal fate acquisition. We have also shown that stimulation with WNT3A+ACTIVIN results in the formation of a human organizer that can induce a secondary axis when grafted into chick embryo^12^, and that the control of the WNT based patterning is primarily due to boundary forces transduced by E-CAD and WNT induced negative feedback from DKK1.

In this work we use our gastruloid model to construct a rudimentary fate map of the human primitive streak. We find that different subpopulations of endoderm or mesoderm emerge robustly from each set of gastruloids depending on BMP, WNT, and NODAL levels, and that by comparison to the mouse embryo we can arrange these subpopulations along an anterior-posterior axis. We also find that there are robust cell migrations from the primitive streak region of each gastruloid and that the character of these migrations depends on what fates the differentiating cells commit to, with fast single-cell migrations in the case of endoderm for instance, or slower group-migration in the case of mesoderm populations. These migrating cells also appear to involute and move under the epiblast section of each gastruloid and in some conditions this migration correlates with the appearance of a COLLAGEN IV basement membrane separating the two layers, mirroring the COLLAGEN IV basement membrane that separates the epiblast from PS derivatives in the mouse. In the absence of direct characterization in human gastrula, our study assumes that the markers we use for hESCs will demarcate the same relative cell types and territories *in vivo*. We believe that this assumption is safe as regardless of the geometry we have tried to use mutually exclusive sets of markers that always delineate the same cell type across multiple vertebrate species, and that have been used for example in recent successful mappings of mouse gastruloid cell populations to the mouse embryo^13^. Taken together, we believe that our gastruloid model is a robust and rich system through which a rudimentary fate map of the primitive streak and a corresponding picture of early human gastrulation can begin to be pieced together.

## Results and Discussion

### Anterior-posterior fate specification in human gastruloids

Building on our previous characterization of gastruloid cell fates^10^ and incentivized by the discovery of a human organizer in WNT3A+ACTIVIN treated cells^12^, we hypothesized that other primitive streak cell subtypes were present in our gastruloids and that they could also be compared to the anterior-posterior axis of the mouse embryo. Our hypothesis was supported by recent work successfully mapping cell types in mouse gastruloids to mouse embryos^13^. In those studies, specific combinations of cell type specific transcription factors were used to identify discrete fates and compare their pattern with the mouse embryo. Here we follow the same strategy and analyze our BMP4, WNT3A, WNT3A+SB, or WNT3A+ACTIVIN induced human gastruloids for anterior-posterior identity and compare them with the mouse gene map and fate map at E7.5 to shed some light on spatial structure of the human PS (Figure 1A-B).

**Figure 1.**
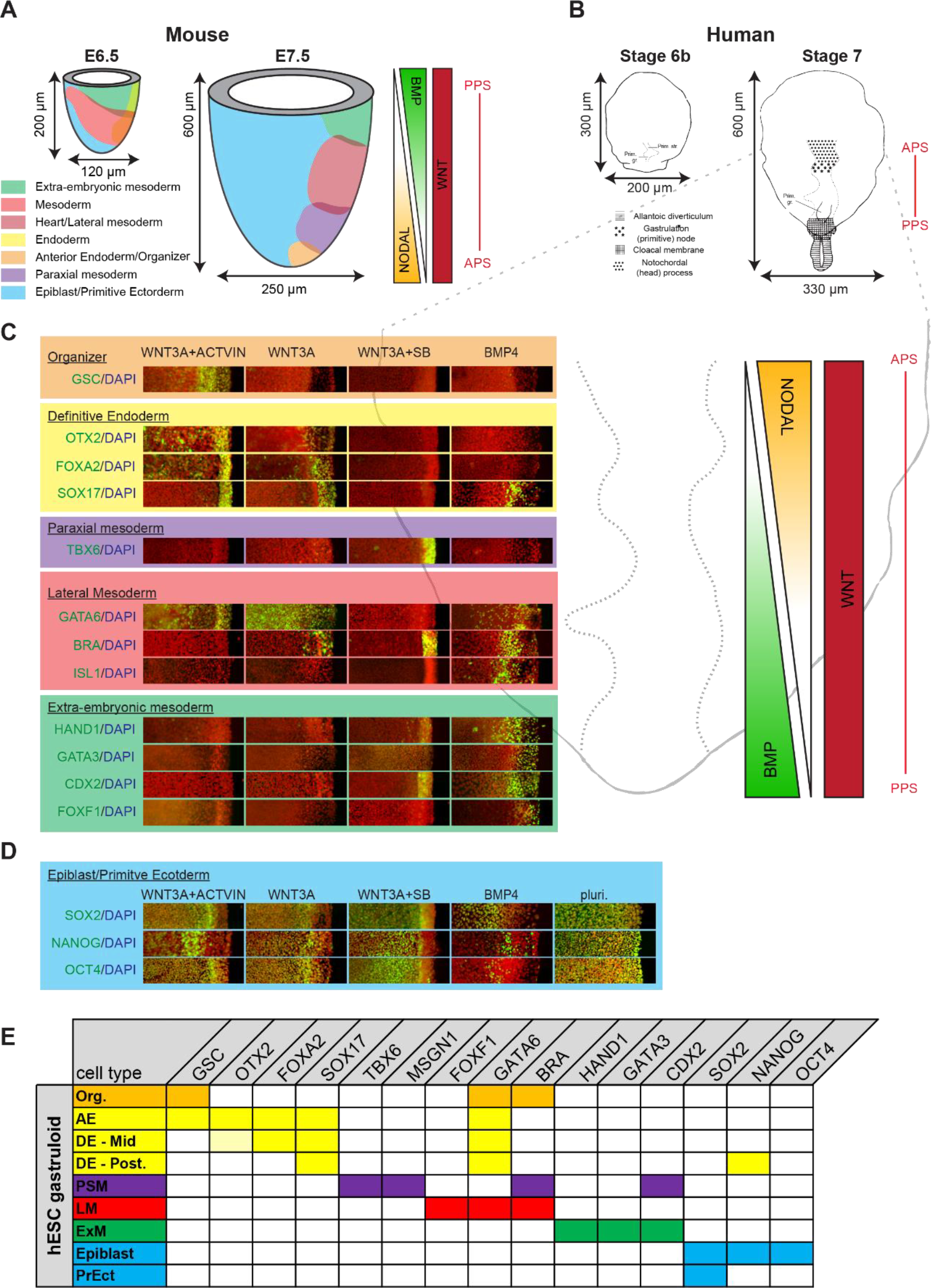
Mapping gastruloid fates to the human primitive streak. (A) Reproduction of the mouse primitive streak fate map from Tam and Behringer 1997. On the right are the inferred signalling gradients of BMP, WNT3A, and Nodal^43,44^. (B) Dorsal graphical representation of the human primitive streak fate map from the Carnegie Collection^45^ (Fig. 6-5. and Fig. 7-4). (C) Mapping of gastruloids stimulated with either BMP4, WNT3A, WNT3A+SB, or WNT3A+Activin to the Carnegie Collection (C.C.) stage 7 human primitive streak. Gastruloids were fixed after 48 h stimulation and stained for the indicated sets of markers. Since each staining is radially symmetric, only a section from r=0 to r=R (500 μm) is shown. (D) Table summarising the expression of each marker for each classified sub-type. The OTX2/DE-Mid box with lighter yellow indicates that expression of OTX2 is less than that observed in other cells in other conditions. Org. = organizer, AE = anterior endoderm, DE-Mid = mid-streak definitive endoderm, DE - Post. = posterior definitive endoderm, PSM = presomitic mesoderm, LM = lateral mesoderm, ExM = extra-embryonic mesoderm, PE = primitive ectoderm.

Strikingly, we found self-organized and largely homogenous anterior-posterior subpopulations that arose distinctly in one set of stimulation conditions and not the others. For instance, only BMP4 induced expression of HAND1, CDX2, and GATA3, and these markers were all present in the same set of cells on the periphery of the gastruloid (Figure 1C). In the mouse HAND1 is first expressed at E7.5 in the trophectoderm and extra-embryonic mesoderm, including the amnion, chorion, allantois and visceral yolk sac^14^. GATA3 is expressed in the mouse and human pre-implantation trophoblast^9,15,16^ and in the mouse E7.5 extra-embryonic ectoderm and allantois^16,17^. CDX2 is also expressed in the mouse and human pre-implantation trophoblast^9^ and in mouse is restricted to the extra-embryonic ectoderm, mesoderm, and posterior endoderm until E8.5^13,18,19^. FOXF1, which beginning at E7.5 in the mouse turns on and marks the lateral plate mesoderm and yolk sac and allantois^20,21^, is also most highly expressed in cells in this region. Based on these comparisons, we take this region of the BMP4 induced gastruloid to most closely resemble the mouse E7.5 extra-embryonic mesoderm. This is further supported by the fact that there is a BMP source from the extra-embryonic ectoderm immediately adjacent to it *in vivo*. The fact that we do not see significant BRA expression in these cells as was found in mouse^13^ may be due to species-specific timing differences (for example we have shown previously that there is a wave of BRA expression earlier in this region at 12-36h^10^). Radially interior to this extra-embryonic mesoderm population are three other readily identifiable subpopulations that are unique to the BMP4 gastruloid. First, in the region adjacent to the extra-embryonic mesoderm there is a population of BRA+/GATA6+/ISL1+ cells. In the mouse at E7.5 GATA6 marks the parietal and definitive endoderm plus the lateral mesoderm^13,22^, while ISL1 first appears at E8.5 and also marks cells in the lateral mesoderm^23,24^. Thus we identify this subpopulation as lateral mesoderm. Second, staining for SOX17 (a marker of definitive endoderm first apparent in mouse at E7-7.5^13^), NANOG (maker of definitive endoderm and epiblast), and OTX2 (marker of anterior epiblast and anterior PS in mouse from E7^13^) detected a population of the SOX17+/NANOG-/OTX2-cells. Based on these markers we identify this population as posterior endoderm. Finally, staining for SOX2 (marker of primitive ectoderm), and OCT4 (marker of epiblast) revealed a SOX2+/NANOG-/OCT4-subpopulation indicative of primitive ectoderm. Together, these four subpopulations in the BMP4 stimulated gastruloid all approximately match the E7.5 proximal posterior primitive streak in mouse.

With WNT3A+SB stimulation we found selective expression of TBX6 in the region that co-expresses CDX2 and BRA (Figure 1C). TBX6 did not appear in the other stimulation conditions, and using qPCR we also found that MSGN1 was selectively induced with WNT3A+SB only (Figure S1A). In the mouse, both TBX6 and MSGN1 are first expressed in the primitive streak in the same region as BRA at E7.5, only to become restricted to the paraxial mesoderm by E8.5^25–27^. The fact that we do not detect significant TBX6 or MSGN1 levels at earlier times in any of the other gastruloids where we also see BRA (data not shown) may reflect a species specific difference between human and mouse. Additionally, although we use CDX2 in our panel of markers for the BMP4 induced gastruloids, CDX2 has also been shown to be critical for paraxial mesoderm development in the mouse and is detectable there from E8.5 onwards^18,28^. The union of these three molecular markers is thus highly suggestive of paraxial mesoderm, and a corresponding time of ~E7.5-8.5 in the mouse.

In the case of WNT3A and WNT3A+ACTIVIN, stimulation led to selective co-expression of the transcription factors FOXA2 and OTX2 in the SOX17+ region at the edge (Figure 1C). In the mouse FOXA2 begins to be expressed in the anterior primitive streak at E7, and becomes restricted to the anterior definitive endoderm and axial mesoderm by E7.75^13,29^. Thus the FOXA2+/OTX2+/SOX17+/BRA-provides the signature of anterior endoderm. Additionally, at 24 hours with WNT3A+ACTIVIN, but not WNT3A alone, we can detect the organizer marker GSC. As previously shown^12^ we can identify an organizer population at 24 h in WNT3A+ACTIVIN gastruloids when GSC is co-expressed with BRA and SOX17 is not yet visible. Finally, the centers of the WNT3A, WNT3A+Activin, and WNT3A+SB stimulated gastruloids differ from the center region of the BMP4 stimulated gastruloids in that they still express NANOG and OCT4, albeit at a lower level than in pluripotency (Figure 1C). We thus categorize these regions as epiblast and not as primitive ectoderm.

A summary of all of the readily identifiable subpopulations is provided in Figure 1E, and a direct comparison with the mouse embryo with references for the expression pattern for each marker is given in Supplementary Table 1. Overall, we find good agreement between the mouse embryo and the gastruloid subpopulations.

### Cell migration

In addition to the emergence of distinct mesoderm and endoderm subtypes in different anterior-posterior positions along the primitive streak, vertebrate gastrulation is also characterized by highly orchestrated cell migrations through the streak and under the epiblast. Indeed, fate specification and migration occur concomitantly.

To track cells in our gastruloid system, we used the ePiggybac transposable element system^30^ to derive clonal RUES2-KiKGR-RFP657-H2B cell lines that contain the photo-convertible protein KikGR and the far-red histone localized fluorescent protein RFP657-H2B. KikGR protein normally fluoresces in green but permanently converts to red upon UV excitation. KikGR also has a long life-time, enabling the detection of cells in which the protein has been switched to the red state even after two days. This tool allows photo-conversion of cells in specific regions of gastruloids and determination of their location after a window of time. More specifically, we used a digital micro-mirror to direct a 405 nm laser to illuminate one of three different annular regions: A_1_, all cells <50 μm from the colony center; A_2_, all cells in a ring >200 μm and <250 μm from the colony center; and A_3_, all cells >400 μm from colony center (Figure 3B-C). Immediately after photo-conversion, micropatterns were stimulated with either control medium, BMP4, WNT3A, WNT3A+ACTIVIN or WNT3A+SB, and imaged to establish the starting point. The same colonies were imaged again at 24 h (Figure S2A), and again at 52 h (Figure 2C and quantified in D).

**Figure 2.**
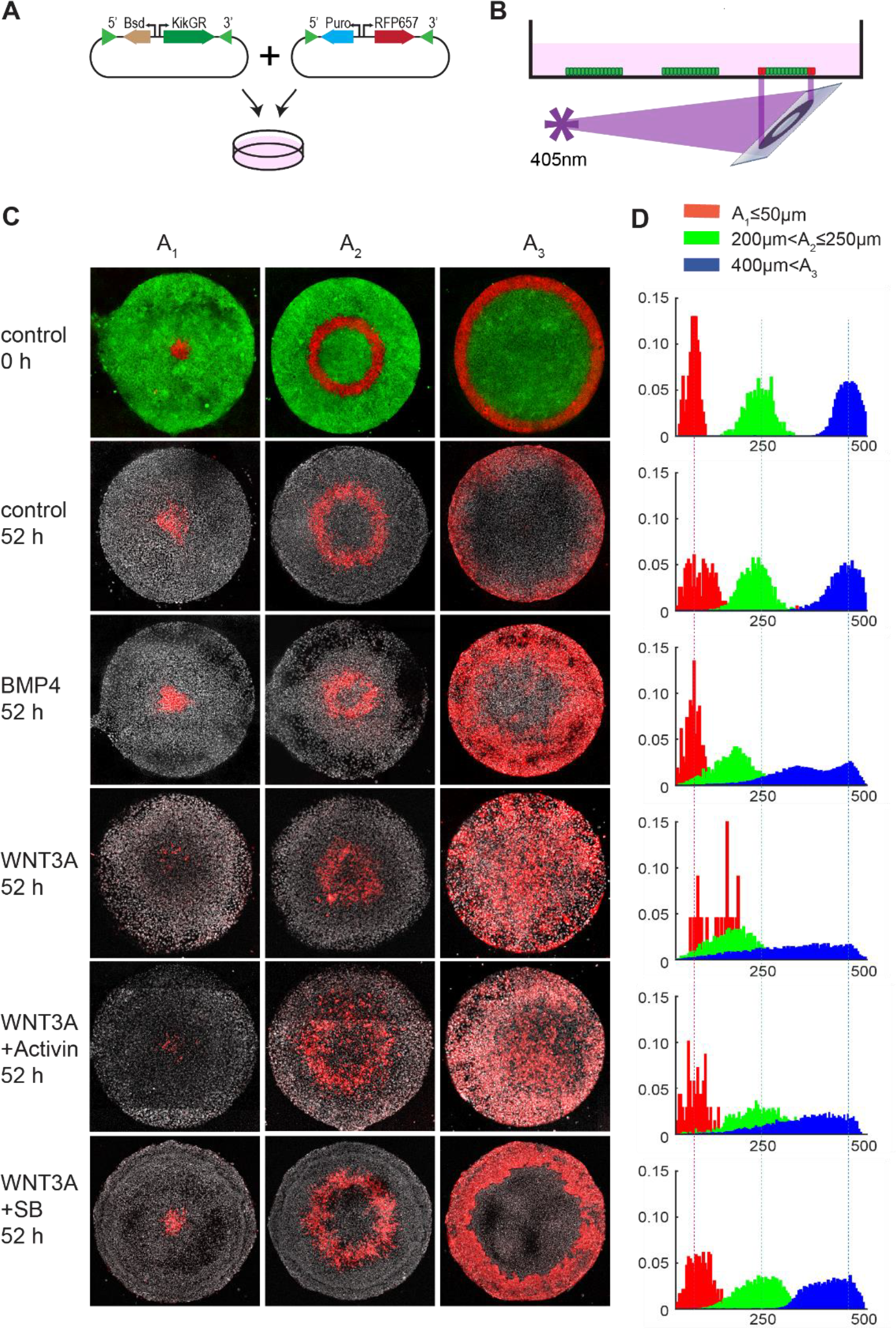
Directed cell migration in the PS region. (A) Cloning strategy to make the RUES2-KiKGR-RFP657-H2B cell line. (B) Using a digital micromirror, annular regions of micropatterned RUES2-KiKGR-RFP657-H2B colonies were selectively exposed to 405 nm light for 3 seconds and permanently switched from green to red fluorescence. (C) Three different annular regions were photoconverted: A_1_ (<50 μm from colony center), A_2_ (>200 μm and <250 μm from colony center), and A_3_ (>400 μm from colony center). After photo-conversion cells were stimulated with either WNT3A, WNT3A+Activin, WNT3A+SB, BMP, or blank media and imaged at 0 h, 24 h (Supplemental Figure 2), and 52 h. First row shows unconverted KikGR fluorescence (green) and converted KikGR fluorescence (red) at 0 h. All other rows show just converted KikGR fluorescence (red) and the far-red histone nuclear marker (grey) at 52 h. In all conditions significant movement of cells in the A_3_ region is observed. (D) Quantification of C (see Methods).

**Figure 3.**
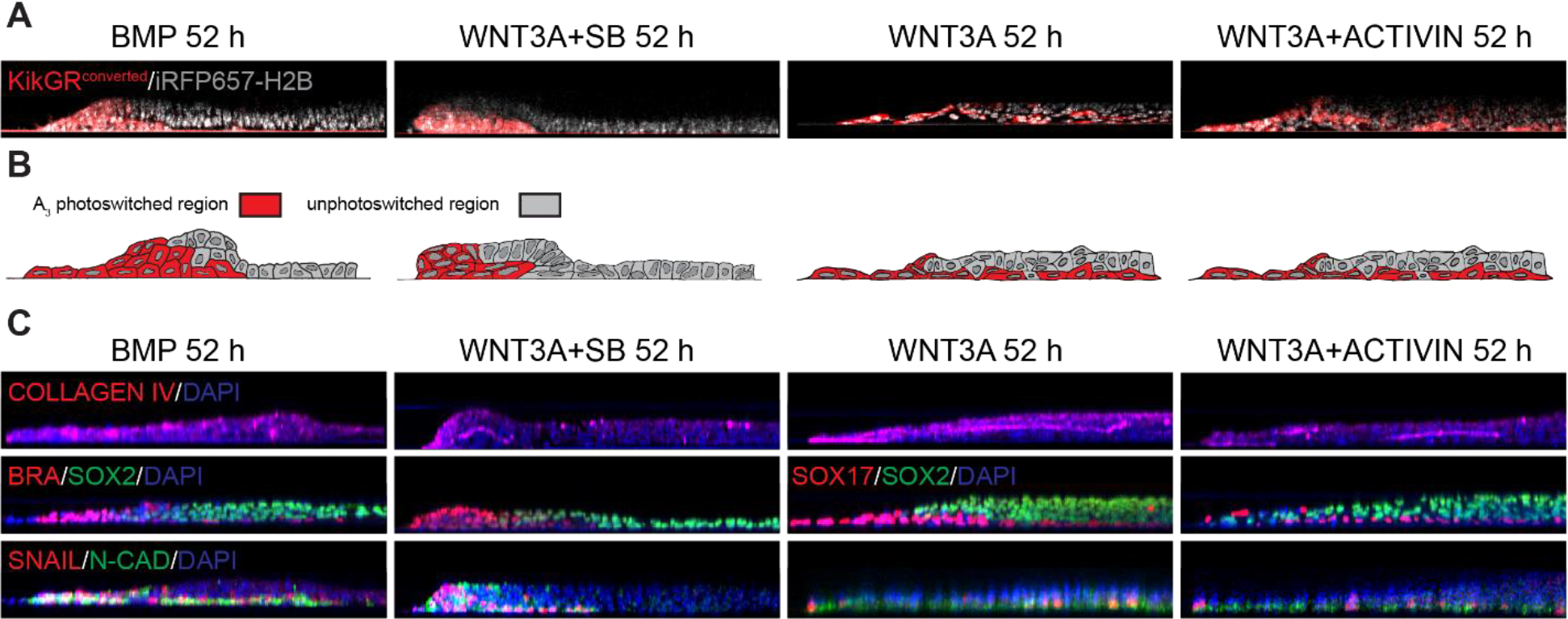
3D gastruloid morphology. (A) Radial cross-sections of RUES2-KiKGR-RFP657-H2B micropatterns photoconverted in region A_3_ and stimulated with WNT3A, WNT3A+Activin, WNT3A+SB, or BMP4 for 52 h. In all conditions the photoconverted (red) cells can be seen to be migrating under the inner epiblast or ectoderm-like region. (B) Hand-drawn depiction of 3D structure of gastruloids inferred from (A). (C) Radial cross-sections of RUES2 gastruloids stimulated with WNT3A, WNT3A+Activin, WNT3A+SB, or BMP4 and fixed and stained at 52 h for the indicated markers. As can be seen by comparison with (A) and (B), the migratory cells are also differentiated to either mesoderm or endoderm and express PS markers. In all the conditions except BMP4, one can also see that a basement layer of COLLAGEN IV separates the migrating cells from the undifferentiated epiblast-like cells overtop, as would be expected in vivo.

We found that in the unstimulated micropatterns the photoconverted cells retained their original position even after 52 h (Figure 2C, 1^st^ and 2^nd^ row). Stimulation with BMP4, WNT3A, WNT3A+ACTIVIN, or WNT3A+SB, however, led to migration of cells localized at the edge (A_3_) towards the center (Figure 2C, rows 3-6), and the onset of these migrations correlated well with the previously reported^12^ epithelial-to-mesenchymal transition (EMT) onset observed in each condition (Figure S2). Under WNT3A and WNT+ACTIVIN stimulation migration started shortly after 24 hours of stimulation and cells migrated in a dispersed, individual manner, travelling long distances between the edge of the colony to the center (Figure 2C, 4^th^ and 5^th^ rows, and Figure S2). In contrast, a slower, shorter, and more compact migration was observed in the BMP4 and WNT3A+SB induced gastruloids (Figure 2C, 3^rd^ and 6^th^ rows). Quantification of the photoconverted cells in the BMP4 treatment revealed two distinct populations: one that remained on the outer edge, and another that migrated inwards (Figure 2C and D, 3^rd^ row). Finally, while no migration was observed in A_1_ regardless of the stimulation, cells in the A_2_ region shifted slightly inward by 52 h following WNT3A+ACTIVIN, WNT3A and BMP4 stimulation. However, these cells do not express EMT markers early on^12^, and it is hard to differentiate between active movement and passive movement as the result of being pushed in by the migration of cells from A_3._ For instance, we speculate that as the A_2_ region in BMP4 gastruloids is more compact than the WNT treated gastruloids this is more the result of pushing from the exterior cells rather than autonomous movement.

To better understand how the cells migrate in each of the conditions we also examined the 3D structure of the gastruloids and what the corresponding fate markers of the migrating cells are. In the WNT3A and WNT3A+ACTIVIN gastruloids the migrating cells express SOX17 and so belong to the Anterior DE subpopulation identified previously. In the BMP4 gastruloids the migratory cells express BRA and so mostly belong to the LM subpopulation. In the WNT3A+SB gastruloids the migratory cells also express BRA and so are the PSM fated cells (Figure 3A and B). In all cases the migrating cells appear to push under the inner epiblast section en route towards the center of the gastruloid (Figure 3A). The nature of this attachment and the interaction of these cells with the migratory cells is also related to the COLLAGEN IV layer that we detect separating these layers in the WNT3A, WNT3A+ACTIVIN, and WNT3A+SB gastruloids (Figure 3C). In the mouse embryo the formation of a COLLAGEN IV basement membrane precedes gastrulation, but here it is unclear if the layer also exists prior to stimulation, or it is produced from one or both populations of cells as differentiation proceeds.

The fact that the observed cell migrations are robust, concurrent with EMT, and dependent on the fate the cells adopt, suggests that we are seeing movements attempting to fulfill the *in vivo* human gastrulation program. In support of this is the fact that in the mouse mesoderm first migrates as compact “wings”^31,32^ behind a leading edge of more dispersed definitive endoderm fated cells^33,34^, since this is consistent with the rates and behavior of mesoderm and endoderm migrating cells in our gastruloids. Compared with cell migration in the avian primitive streak^35– 39^ not much else is known about mammalian primitive streak cell migration or the mechanisms and chemical cues behind it^40^. We believe that our gastruloid model offers a glimpse of this difficult to study *in vivo* process, and moving forward may present a fruitful alternative approach to dissect the molecular mechanisms underlying cell-migration during a pivotal time of human gastrulation.

### Mapping cell migrations and fates to the human primitive streak

Putting together our gene maps and anterior-posterior signatures, our cell migration patterns, and 3D cross-sections, we are able to suggest a detailed graphical representation at what gastrulation may look like in human PS at various anterior-posterior positions (Figure 4). We propose that the edges of the Epibalst/PrEct region of each gastruloid correspond to the median of the PS, while the centers of each gastruloid are positioned laterally relative to this median. In this schema the direction of migration of differentiating cells (indicated by arrows) is from the medial line of the streak out laterally, underneath the COLLAGEN IV and epiblast or primitive ectoderm layers. The uncovered region of differentiated cells on the edge of our gastruloids we believe would be covered in the embryo since in that anterior-posterior polarized streak geometry we would expect the Epibalst/PrEct region to grow (as in mouse) or flow (as in chick) to fill in that space. Interestingly, whether the migrating cells go under or over appears to be surface dependent, as in previously published work on PDMS micropatterns the corresponding migratory population appeared on the top of the epiblast^12^. We speculate that in both conditions they may be responding to similar cues but taking whichever route is easier depending on attachment of the remaining epiblast/PrEct region to the surface.

**Figure 4.**
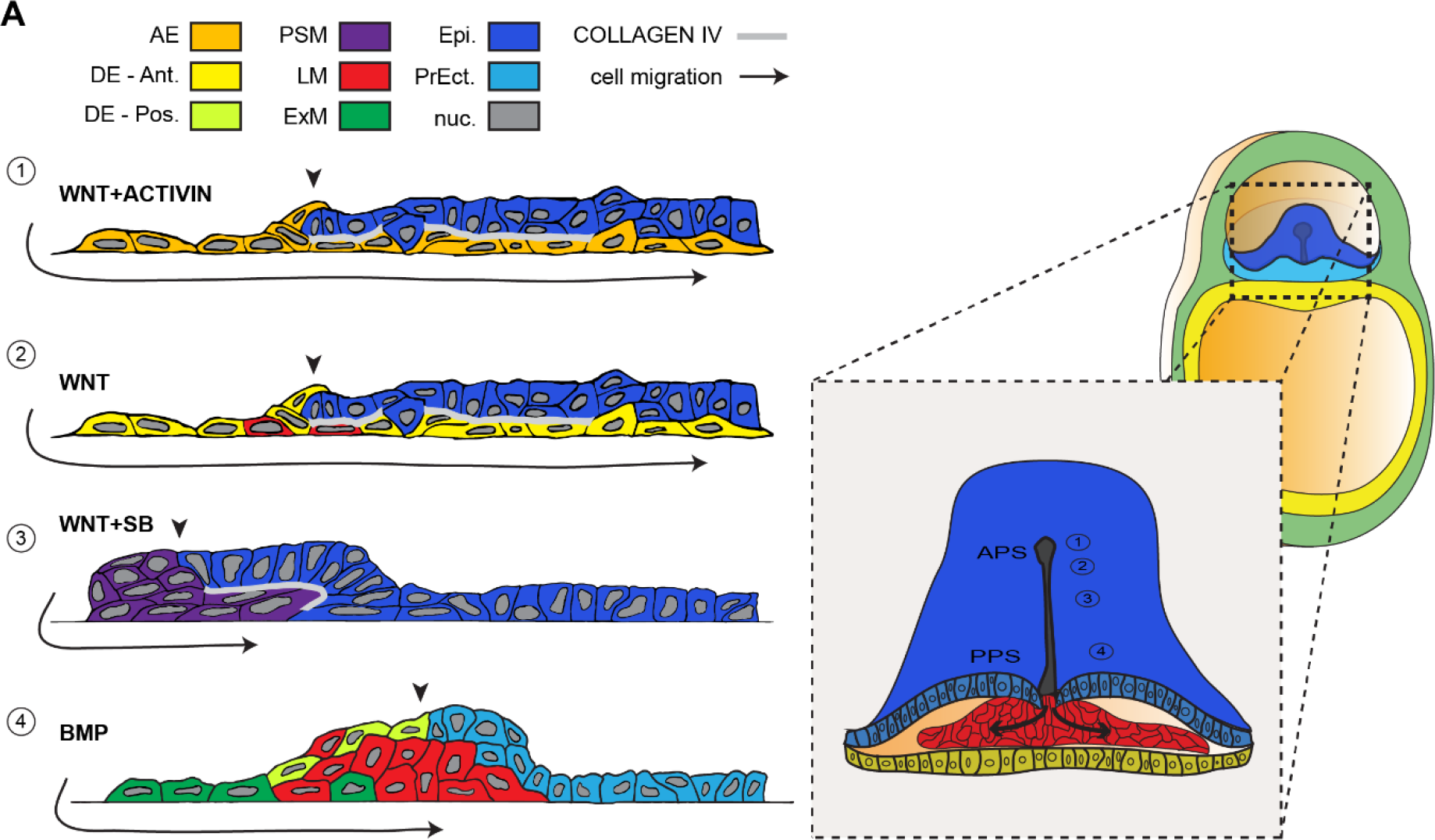
Mapping gastruloid cell migrations and fates to the human primitive streak. (A) Cartoons summarizing the fates and 3D structure of each type of gastruloid at 52 h and mapping to the human embryo. As indicated by the arrowheads in the diagram, we believe the edge of the Epiblast/PrEct region in each gastruloid corresponds to the medial part of the *in vivo* primitive streak, and that our migrations (indicated by arrows) therefore occur medially to laterally.

There is no doubt that our gastruloid derived gene/fate map lacks details and features that could be observed in the developing *in vivo* human embryo. We anticipate that missing cell types, such as germ cells or intermediate mesoderm, for example, might be revealed in the future with the use of single cell RNA-seq of gastruloids and sets of markers informed by new efforts to acquire single cell RNA-seq data from gastrulating primate embryos^41^. There is also the limitation that, unlike the *in vivo* case, our anterior-posterior streak is a composite of separate differently stimulated gastruloids. That said, given what we have learned about the required stimulation conditions for each fate subpopulation, it may be possible with advances in micropatterning techniques or localized ligand sources to recreate the entire anterior-posterior streak in a single micropattern. This would be a superior model and allow much better understanding of the relative timing of EMT, fate specification, and migrations for example. Regardless of the limitations of our current studies, however, we believe our results represent a first step forwards to observing and mapping the origin of fate during our own human development.

## Acknowledgements

We thank Anna-Katerina Hadjantonakis for her gift of pCAG:KikGR. We also thank members of her laboratory and members of the A.H.B. and E.D.S. laboratories for helpful scientific discussions. We especially thank Pablo Ariel and the staff of the Rockefeller Bio-Imaging Resource Center for support in imaging and photoswitching experiments. This work was supported by grants R01 HD080699 and R01 GM101653.

## Materials and Methods

### Cell Culture

hESCs (RUES2 cell line) were grown and maintained in HUESM medium conditioned by mouse embryonic fibroblasts (MEF-CM) and supplemented with 20ng/mL bFGF. Testing for mycoplasma was carried out before beginning each set of experiments and again at 2-month intervals. For maintenance conditions, cells were grown on GelTrex (Invitrogen) coated tissue culture dishes (BD Biosciences, 1:40 dilution). The dishes were coated overnight at 4°C and then incubated at 37°C for at least 10 minutes before the cells were seeded on the surface. Cells were passaged using Gentle Cell Dissociation Reagent (Stem Cell Technologies 07174).

### Micropatterned Cell Culture

We used micropatterned glass coverslips from CYTOO. The coverslips were first coated with 10 ug/ml laminin 521 (Biolamina) diluted in PBS with calcium and magnesium (PBS++) for 3 h at 37°C. Cells were dissociated with StemPro Accutase (Life Technologies) for 7 minutes. Cells were then washed once with growth media, washed again with PBS, and then re-suspended in growth media with 10μM ROCK-inhibitor Y-27632 (Abcam) in 35mm tissue culture plastic dishes. For each coverslip 1×10^6^ cells in 2mL of media were used. After 1 h ROCK-inhibitor was removed and was replaced with standard growth media supplemented with Pen-Strep (Life Technologies). Cells were stimulated with the following ligands or small molecules 12 h after seeding: 100ng/mL WNT3A, 50ng/mL BMP4, 100ng/mL Activin-A, 10μM SB, or 2μM IWP2.

### Establishment of stable photo-convertible hESC cell Line

pCAG:KikGR was a gift from Anna-Katerina Hadjantonakis (Addgene plasmid # 32608). The KikGR protein from this plasmid was amplified with forward primer 5-ATTGGATCCCGGATGGTGAGTGTGATTACATCAGAA-3 and reverse primer 5-TATGCGGCCGCCGGTTACTTGGCCAGCCTTG-3 and, using BamHI and NotI cloning sites, was inserted into an ePiggyBac plasmid with a pCAG promoter and puromycin resistance cassette (Lacoste et al., 2009). This plasmid, along with a plasmid carrying the piggybac transposase and another ePiggyBac plasmid carrying a H2B-RFP657 fluorescent protein and blasticidin resistance, were nucleofected into 1×10^6^ pluripotent RUES2 cells using the B-016 setting on an Amaxa Nucleofector II (Lonza). Nucleofected cells were then plated as per maintenance conditions, but supplemented with 10uM ROCK-inhibitor. Selection for both puromycin and blasticidin commenced after 2 days, and ROCK-inhibitor was maintained until colonies reached adequate size (typically 8-16 cells per colony). To derive pure clones, individual colonies were picked in an IVF hood with a 20 μL pipette tip and seeded into separate wells with growth media and ROCK-inhibitor. Once successfully established, each clone was assayed functionally for brightness and homogeneity of the KikGR and H2B-RFP647 fluorescent proteins. Each clone was also assayed functionally for its ability to recapitulate the self-organization in micropatterns when stimulated with BMP4. Three successful clones were selected, and one was used for subsequent studies.

### Immunostaining

Cultures were fixed in 4% paraformaldehyde for 20 minutes, washed twice with PBS, and then blocked and permeabilized with 3% donkey serum and 0.1% Triton X-100 in PBS for 30 minutes. Cultures were incubated overnight with primary antibodies in this blocking buffer at 4°C (for primary antibodies and dilutions, see Table S1), washed 3 times with PBS+0.1% Tween-20 for 30 minutes each, and then incubated with secondary donkey antibodies (Alexa 488, Alexa 555, Alexa 647) and DAPI for 30 minutes before a final washing with PBS and mounting onto glass slides for imaging.

### qPCR data

RNA was collected in Trizol at indicated time points from either mircopatterned colonies or from small un-patterned colonies and was purified using the RNeasy mini kit (Qiagen). qPCR was performed as described previously^42^ and primer designs are listed in the Table S2.

### Imaging and Image Analysis

Images were acquired with a Zeiss Axio Observer and a 20x/0.8 numerical aperture (NA) lens or with a Leica SP8 inverted confocal microscope with a 40×/1.1-NA water-immersion objective. Image analysis and stitching was performed with ImageJ and custom Matlab routines. For tracking cells with the RUES2-KikGR-RFP657-H2B cell line, segmentation was carried as for fixed cells, except here we used the H2B-RFP647 fluorescence signal instead of a DAPI signal as the nuclear marker. We then trained an Ilastick classifier to binarize cells as photo-converted or unconverted, and binned the converted cells into a radial histogram. The plots in Figure 3D represent the average of n=5 colonies.

### Cell-tracking with photo-convertible line

RUES2-KikGR-RFP657-H2B cells were plated onto micropatterned CYTOO chips instead of home-made chips in order to accommodate the 19.5×19.5mm spaced CYTOO chip holder. Immediately after stimulation with BMP, WNT3A, or WNT3A+SB, each chip was sequentially loaded into the CYTOO chip holder, placed on the microscope, photo-converted, washed with PBS, and then returned to the culture dish. Photo-conversion was carried out on a custom built spinning-disk confocal Inverted Zeiss Axiovert 200 microscope with a Photonics Instruments Digital Mosaic system with a 405nm laser. Regions of Interest (ROIs) for photo-conversion were programmed with custom Matlab code, and then loaded into the Metamorph software used to operate the microscope. Individual colonies were found, aligned with the ROI, and had their stage position stored. Using a custom written Metamorph script, each colony was sequentially imaged with GFP and RFP filters, exposed to 3162ms of 405nm light from the laser, and then imaged again to check for complete photo-conversion. Once photo-converted, each CYTOO chip was returned to its native 35mm dish and placed in an incubator. For tracking these cells at later times, the micropatterned chips were taken out of the incubator and sequentially re-loaded in the CYTOO holder and imaged with the afore-mentioned Lecia SP8 confocal microscope. They were then washed and returned to the incubator.

---

**Supplemental Table S1:** Antibody information

| Antigen | Antibody | Dilution |
| --- | --- | --- |
| BRACHYURY | R&D Systems AF-2085 | 1:300 |
| CDX2 | Abcam Ab-15258 | 1:50 |
| COLLAGEN IV | Abcam Ab-6586 | 1:100 |
| FOXA2 | SCBT 6554 | 1:200 |
| FOXF1 | R&D Systems AF-4798 | 1:200 |
| GOOSECOID | R&D AF-4086 | 1:100 |
| HAND1 | R&D Systems AF-3168 | 1:100 |
| ISL1 | Abcam Ab-109517 | 1:200 |
| NANOG | R&D Systems AF-1997 | 1:200 |
| N-CADHERIN | BioLegend 350802 | 1:200 |
| OCT4 | BD 611203 | 1:400 |
| OTX2 | SCBT 30659 | 1:200 |
| SNAIL | R&D Systems AF-3639 | 1:200 |
| SOX17 | R&D Systems AF-1924 | 1:200 |
| SOX2 | Cell Signalling 3579 | 1:200 |
| TBX6 | R&D Systems AF-4744 | 1:100 |
| GATA3 | Thermofisher MA1-028 | 1:100 |
| GATA6 | Cell Signalling 5851 | 1:400 |

**Supplemental Table S2:**
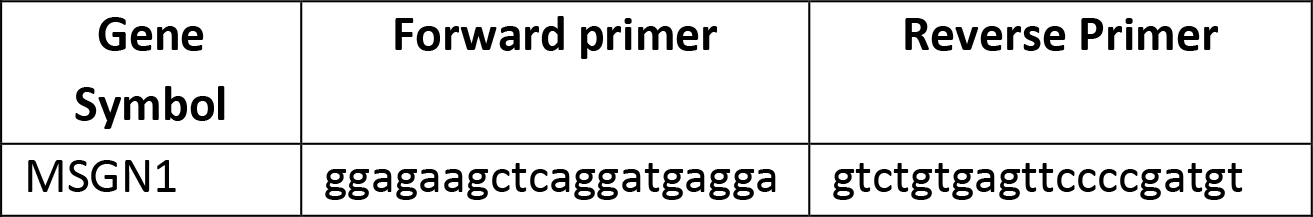
RT-QPCR Primer designs

**Supplemental Figure 1.**
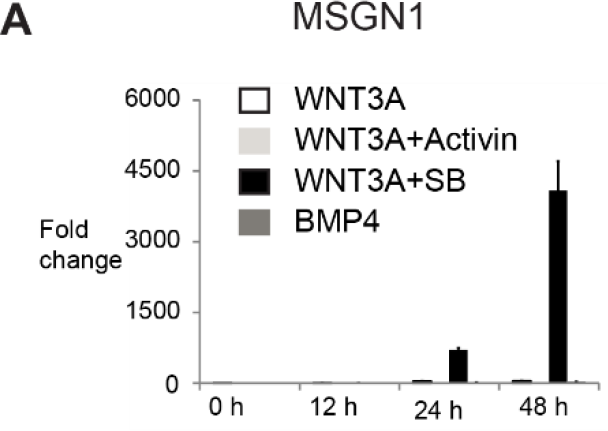
Additional markers of early endoderm and mesoderm subtypes. (A) qPCR for paraxial mesoderm marker MSGN1 shows that it is most highly expressed in WNT3A+SB treated micropatterns at 48 h. Error bars represent the standard deviation of three technical replicates for each condition.

**Supplemental Figure 2.**
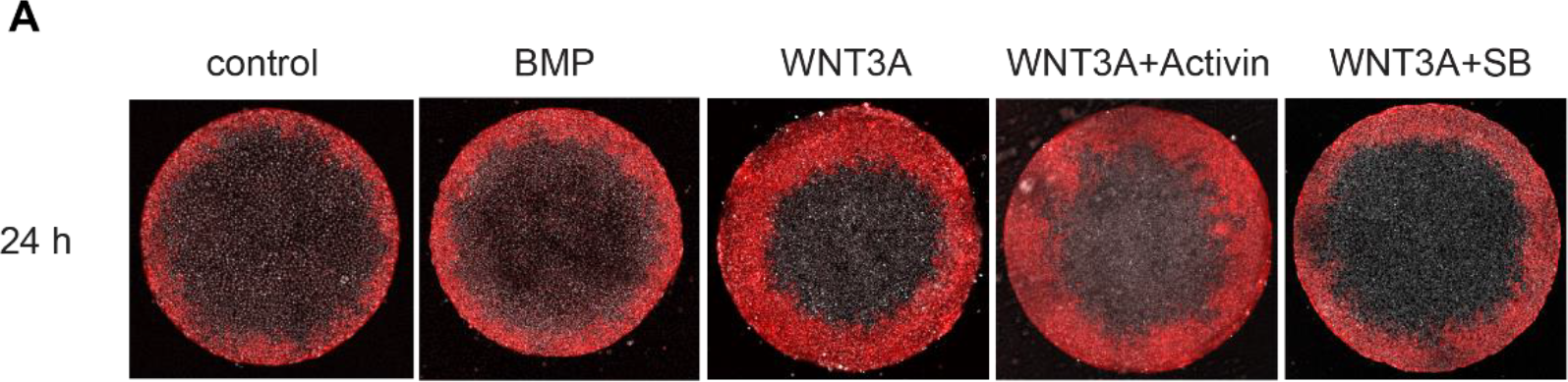
Onset of directed cell migration. (A) Micropatterns with photoconverted cells in region A_3_ (>400 μm from colony center) and stimulated with either WNT3A, WNT3A+Activin, WNT3A+SB, BMP, or blank media were imaged at 24 h. As can be seen by comparison with the blank stimulated colony, cells in the outer region of the WNT3A and WNT3A+Activin micropatterns have started moving inwards at this time, ahead of the corresponding cells in the BMP4 and WNT3A+SB micropatterns.

## References

1. Solnica-Krezel, L. & Sepich, D. S. Gastrulation: Making and Shaping Germ Layers. Annu. Rev. Cell Dev. Biol. 28, 687–717 (2012).

2. Tam, P. P. L. & Behringer, R. R. Mouse gastrulation:the formation of the mammalian body plan. Mech. Dev. 68, 3–25 (1997).

3. Hatada, Y. & Stern, C. D. A fate map of the epiblast of the early chick embryo. Development 120, 2879–89 (1994).

4. Conklin, E. G. The organization and cell lineage of the ascidian egg. J. Acad. Nat. Sci. Philadelphi. 13, 1–119 (1905).

5. Vogt, W. Gestaltungsanalyse am Amphibienkeim mit Örtlicher Vitalfärbung. Wilhelm Roux. Arch. Entwickl. Mech. Org. 120, 384–706 (1929).

6. Kimmel, C. B., Warga, R. M. & Schilling, T. F. Origin and organization of the zebrafish fate map. Development 108, 581–594 (1990).

7. Hyun, I., Wilkerson, A. & Johnston, J. Embryology policy: Revisit the 14-day rule. Nature 533, 169–171 (2016).

8. International Society of Stem Cell Research. Guidelines for Stem Cell Research and Clinical Translation. International Society for Stem Cell Research (2016).

9. Deglincerti, A. et al. Self-organization of the in vitro attached human embryo. Nature 533, 251–254 (2016).

10. Warmflash, A., Sorre, B., Etoc, F., Siggia, E. D. & Brivanlou, A. H. A method to recapitulate early embryonic spatial patterning in human embryonic stem cells. Nat. Methods 11, 847–54 (2014).

11. Simunovic, M. & Brivanlou, A. H. Embryoids, organoids and gastruloids: new approaches to understanding embryogenesis. Development 144, 976–985 (2017).

12. Martyn, I., Kanno, T. Y., Ruzo, A., Siggia, E. D. & Brivanlou, A. H. Self-organization of a human organizer by combined Wnt and Nodal signalling. Nature 558, 132–135 (2018).

13. Morgani, S. M., Metzger, J. J., Nichols, J., Siggia, E. D. & Hadjantonakis, A.-K. Micropattern differentiation of mouse pluripotent stem cells recapitulates embryo regionalized cell fate patterning. Elife 7, e32839 (2018).

14. Firulli, A. B., McFadden, D. G., Lin, Q., Srivastava, D. & Olson, E. N. Heart and extra-embryonic mesodermal defects in mouse embryos lacking the bHLH transcription factor Hand1. Nat. Genet. 18, 266–270 (1998).

15. Home, P. et al. Genetic redundancy of GATA factors in the extraembryonic trophoblast lineage ensures the progression of preimplantation and postimplantation mammalian development. Development 144, 876–888 (2017).

16. Ralston, A. et al. Gata3 regulates trophoblast development downstream of Tead4 and in parallel to Cdx2. Development 137, 395–403 (2010).

17. Manaia, A., Lemarchandel, V., Klaine, M., Romeo, P. & Godin, I. Lmo2 and GATA-3 associated expression in intraembryonic hemogenic sites. Development 127, 643–653 (2000).

18. Beck, F., Erler, T., Russell, A. & James, R. Expression of Cdx-2 in the Mouse Embryo and Placenta: Possible Role in Patterning of the Extra-Embryonic Membranes. Dev. Dyn. 204, 219–227 (1995).

19. Sherwood, R. I., Maehr, R., Mazzoni, E. O. & Melton, D. A. Wnt signaling specifies and patterns intestinal endoderm. Mech. Dev. 128, 387–400 (2011).

20. Peterson, R. S. et al. The winged helix transcriptional activator HFH-8 is expressed in the mesoderm of the primitive streak stage of mouse embryos and its cellular derivatives. Mech. Dev. 69, 53–69 (1997).

21. Mahlapuu, M., Ormestad, M., Enerback, S. & Carlsson, P. The forkhead transcription factor Foxf1 is required for differentiation of extra-embryonic and lateral plate mesoderm. Development 128, 155–166 (2001).

22. Koutsourakis, M., Langeveld, A., Patient, R., Beddington, R. & Grosveld, F. The transcription factor GATA6 is essential for early extraembryonic development. Development 126, 723–732 (1999).

23. Cai, C.-L. et al. Isl1 identifies a cardiac progenitor population that proliferates prior to differentiation and contributes a majority of cells to the heart. Dev. Cell 5, 877–889 (2003).

24. Zhuang, S. et al. Expression of Isl1 during mouse development. Gene Expr. Patterns 13, 407–412 (2013).

25. Chapman, D. L., Agulnik, I., Hancock, S., Silver, L. M. & Papaioannou, V. E. Tbx6, a mouse T-Box gene implicated in paraxial mesoderm formation at gastrulation. Dev. Biol. 180, 534–542 (1996).

26. Yoon, J. K., Moon, R. T. & Wold, B. The bHLH class protein pMesogenin1 can specify paraxial mesoderm phenotypes. Dev. Biol. 222, 376–391 (2000).

27. Chalamalasetty, R. B. et al. Mesogenin 1 is a master regulator of paraxial presomitic mesoderm differentiation. Development 141, 4285–4297 (2014).

28. Savory, J. G. A. et al. Cdx2 regulation of posterior development through non-Hox targets. Development 136, 4099–4110 (2009).

29. Nowotschin, S. et al. Charting the emergent organotypic landscape of the mammalian gut endoderm at single-cell resolution. bioRxiv 471078 (2018). doi:10.1101/471078

30. Lacoste, A., Berenshteyn, F. & Brivanlou, A. H. Resource An Efficient and Reversible Transposable System for Gene Delivery and Lineage-Specific Differentiation in Human Embryonic Stem Cells. Stem Cell 5, 332–342 (2009).

31. Parameswaran, M. & Tam, P. P. Regionalisation of cell fate and morphogenetic movement of the mesoderm during mouse gastrulation. Dev. Genet. 17, 16–28 (1995).

32. Sutherland, A. E. Tissue morphodynamics shaping the early mouse embryo. Semin. Cell Dev. Biol. 55, 89–98 (2016).

33. Viotti, M., Nowotschin, S. & Hadjantonakis, A.-K. SOX17 links gut endoderm morphogenesis and germ layer segregation. Nat. Cell Biol. 16, 1146–1156 (2014).

34. Rivera-Perez, J. A. & Hadjantonakis, A.-K. The Dynamics of Morphogenesis in the Early Mouse Embryo. Cold Spring Harb. Perspect. Biol. 7, (2014).

35. Hardy, K. M. et al. Non-canonical Wnt signaling through Wnt5a/b and a novel Wnt11 gene, Wnt11b, regulates cell migration during avian gastrulation. Dev. Biol. 320, 391–401 (2008).

36. Yang, X., Chrisman, H. & Weijer, C. J. PDGF signalling controls the migration of mesoderm cells during chick gastrulation by regulating N-cadherin expression. Development 135, 3521–3530 (2008).

37. Yang, X., Dormann, D., Munsterberg, A. E. & Weijer, C. J. Cell movement patterns during gastrulation in the chick are controlled by positive and negative chemotaxis mediated by FGF4 and FGF8. Dev. Cell 3, 425–437 (2002).

38. Yue, Q., Wagstaff, L., Yang, X., Weijer, C. & Munsterberg, A. Wnt3a-mediated chemorepulsion controls movement patterns of cardiac progenitors and requires RhoA function. Development 135, 1029–1037 (2008).

39. Sweetman, D., Wagstaff, L., Cooper, O., Weijer, C. & Munsterberg, A. The migration of paraxial and lateral plate mesoderm cells emerging from the late primitive streak is controlled by different Wnt signals. BMC Dev. Biol. 8, 63 (2008).

40. Stankova, V., Tsikolia, N. & Viebahn, C. Rho kinase activity controls directional cell movements during primitive streak formation in the rabbit embryo. Development 142, 92–98 (2015).

41. Nakamura, T. et al. Single-cell transcriptome of early embryos and cultured embryonic stem cells of cynomolgus monkeys. Sci. data 4, 170067 (2017).

42. Etoc, F. et al. A Balance between Secreted Inhibitors and Edge Sensing Controls Gastruloid Self-Organization. Dev. Cell 39, 302–315 (2016).

43. Tam, P. P. L. & Loebel, D. a F. Gene function in mouse embryogenesis: get set for gastrulation. Nat. Rev. Genet. 8, 368–81 (2007).

44. Arnold, S. J. & Robertson, E. J. Making a commitment: cell lineage allocation and axis patterning in the early mouse embryo. Nat. Rev. Mol. Cell Biol. 10, 91–103 (2009).

45. O’Rahilly, R. & Müller, F. Developmental Stages in Human Embryos. (Carnegie Institution of Washington, 1987).

